# Loudness and sound category: Their distinct roles in shaping perceptual and physiological responses to soundscapes

**DOI:** 10.1101/2025.04.04.647310

**Authors:** Mercede Erfanian, Tin Oberman, Maria Chait, Jian Kang

**Author notes:** **Corresponding author:** Mercede Erfanian.

## Abstract

When compared to nature sounds, exposure to mechanical sounds evokes higher levels of perceptual and physiological arousal, prompting the recruitment of attentional and physiological resources to elicit adaptive responses. However, it is unclear whether these attributes are solely related to the sound intensity of mechanical sounds, since in most real-world scenarios, mechanical sounds are present at high intensities, or if other acoustic or semantic factors are also at play. We measured the Skin Conductance Response (SCR), reflecting sympathetic nervous system (SNS) activity as well as the pleasantness and eventfulness of the soundscape across two passive and active listening tasks in (N = 25) healthy subjects. The auditory stimuli were divided into two categories, nature, and mechanical sounds, and were manipulated to vary in three perceived loudness levels. As expected, we found that the sound category influenced perceived soundscape pleasantness and eventfulness. SCR was analysed by taking the mean level across the stimulus epoch, and also by quantifying its dynamic. We found that mean SCR was modulated by loudness only. SCR rise-time (a measure of speed of the skin response) correlated significantly with soundscape pleasantness and eventfulness for nature and mechanical sounds. This study highlights the importance of considering both loudness level and sound category in evaluating the perceptual soundscape, highlighting SCR as a valuable tool for such assessments.

**Impact statement:** Loud mechanical sounds are considered more unpleasant than nature sounds. This study examined whether loudness alone causes this effect or if the type of sound (e.g., mechanical vs. Nature) also plays a role. By measuring skin conductance reflecting the automatic and unconscious activation of the body’s autonomic system, we found that physical reactions are driven by loudness but also correlate with the listener’s judgement of the pleasantness and eventfulness (describing the perceived level of activity or stimulation) of a soundscape.

## Introduction

“Soundscape” is defined as *an acoustic environment perceived, experienced and/or understood by a person or people, in context* (ISO12913-1, 2014). It is proposed to be composed of two main perceptual attributes, pleasantness and eventfulness. These attributes are considered to be distinct from the intrinsic physical characteristics of the acoustic environment and function as evaluative metrics for auditory quality (Erfanian et al., 2021; ISO, 12913-3: 2019). Pleasantness encapsulates the emotional resonance and affective magnitude of auditory perception, while eventfulness encompasses the perceptual intensity and dynamic variability of the auditory experience (Erfanian et al., 2021; Erfanian et al., 2019; Kang et al., 2019).

Considerable evidence suggests that particular acoustic properties, known as primary factors, may influence the soundscape (McDermott, 2012). These factors may include the spectral content of sounds, which contain substantial concentrations of energy in varying frequency ranges (Kumar et al., 2008; Patchett, 1979), or specific temporal modulations (Arnal et al., 2019; Kumar et al., 2008). The composition of these acoustic properties may be inherent to different sound sources (e.g., traffic noise and a public park) in the context of urban areas which gives rise to the variance in the appraisal of those sound sources, making them pleasant or unpleasant (Bradley & Lang, 2000a, 2000b; Gomez & Danuser, 2004; Hume & Ahtamad, 2013; Medvedev et al., 2015). Mounting evidence suggests that nature sounds like ocean waves characterized by certain acoustic properties (high energy in low frequencies), contribute to the psychological and physiological benefits (through increased activity of the Parasympathetic Nervous System (PSNS)) (Alvarsson et al., 2010; Bradley & Lang, 2000a; Buxton et al., 2021; Hedblom et al., 2019; Li & Kang, 2019; Medvedev et al., 2015). In contrast, mechanical sounds like sirens with high energy in high frequencies, induce unpleasantness, accompanied by an increase in the Sympathetic Nervous system (SNS) activity which can be quantified through physiological indicators such as Skin Conductance Response (SCR) (Li & Kang, 2019; Medvedev et al., 2015). Additionally, a primary factor that determines unpleasantness is the loudness level (Carraturo et al., 2024; Mitchell et al., 2021; Oszczapinska et al., 2024; Skagerstrand et al., 2017).

Despite the availability of in-situ studies (Aletta & Torresin, 2023; Axelsson et al., 2010; Erfanian et al., 2021; Jo & Jeon, 2020; Tarlao et al., 2022), there remains a dearth of evidence regarding the nature of unpleasantness assessments in response to mechanical sounds. This issue emerges because, in most urban real-world scenarios, mechanical sounds are usually associated with higher intensity than nature sounds. In addition, previous laboratory-based research presented mechanical sounds as louder relative to nature sounds, eliciting stronger perceptual and physiological representations (Gomez & Danuser, 2004; Hume & Ahtamad, 2013; Li & Kang, 2019; Medvedev et al., 2015; Tavano & Poeppel, 2019). Therefore, it remains inconclusive whether overall loudness alone influences these perceptual and physiological attributes, or if the combination of overall loudness with other unique acoustic features accounts for the observed differences between nature and mechanical sounds.

SCR is a phasic, stimulus-locked change in the electric conductivity of the skin. It has been widely used to measure physiological activity in response to sounds (Benedek & Kaernbach, 2010; Boucsein, 2012; Bradley & Lang, 2000a, 2000b; Dawson et al., 2007; Gatti et al., 2018; Gomez & Danuser, 2004; Greco et al., 2016; Lang & Bradley, 2010; Li & Kang, 2019; Medvedev et al., 2015; Schweiger & Maltzman, 1985). Under consistent environmental conditions (e.g., room temperature), the amplitude of the sudomotor nerve burst, driven by the intensity of stimuli such as sound (SPL), is linearly related to the number of recruited sweat glands and the corresponding SCR (Bach, 2014; Benedek & Kaernbach, 2010; Wallin, 1981). Sweat glands are predominantly innervated by sympathetic cholinergic fibres originating from the sympathetic chain (Shields et al., 1987). Additionally, the activity of sweat glands is strongly modulated by the limbic system, which is involved in affective sound processing (Fruhholz & Grandjean, 2013; Fruhholz et al., 2016). Beyond SCR amplitude, additional indices can be leveraged in experimental paradigms including SCR rise-time. The SCR rise-time is the temporal interval between the onset of the response (SCR initiation) and its peak amplitude, during which the current takes to rise from 10% to 90% of its final value (Boucsein, 2012; Dawson et al., 2007). The rise-time of SCR offers insights into the speed of the physiological response to a stimulus (Dawson et al., 2007; Jindrová et al., 2020; Venables & Christie, 1980). A shorter rise-time indicates a more rapid physiological response, which can be seen in situations where a stimulus elicits a strong emotional response. A longer rise-time may be indicative of a more gradual or muted physiological response, which may occur in response to a weaker or less emotionally salient stimulus. Taken together, this evidence suggests that SCR is a useful method for quantifying the physiological basis of soundscape properties.

Our study aims to address two primary objectives: first, is to determine whether loudness, a percept driven by the sound pressure level (SPL in dB) as well as frequency (in Hz), contributes to variance in the pleasantness and eventfulness and the underlying SCR. The second aim is to examine the effects of two distinct sound categories, nature and mechanical, on pleasantness and eventfulness, and their associated SCR. This classification of sounds is supported by previous research on sound taxonomy (Bones et al., 2018; Salamon et al., 2014).

To address these objectives, we measure the pleasantness, eventfulness, and SCR in response to complex, single-sourced nature and mechanical sound scenarios presented at three loudness levels of low (10 sones), medium (20 sones), and loud (30 sones) over a period of ∼15 s in two separate listening tasks (passive and active) in 25 healthy participants. We expect that loudness would lead to a decrease in perceptual pleasantness, whereas it would result in an increase in eventfulness and SCR. Furthermore, if overall intensity is the only inherent characteristic leading to differences between nature and mechanical sounds, we predict that nature and mechanical sounds at the same loudness level (e.g., 10 sones – low) derive similar subjective and objective responses.

## Methods

### Participants

Thirty-two paid participants took part in this study (17 females; age mean=28.3 ± 10.61, age range 18 - 45). They reported no hearing/auditory difficulties/impairment, and no neurological or relevant health dysfunctions. The same cohort of participants engaged in both the initial passive listening task and the subsequent active listening task. All participants were briefed on the experimental protocol, provided written informed consent, and were remunerated for their participation. The experimental procedures were approved by the Ethics Committee of University College London.

We excluded one participant due to inattentiveness during the passive listening task. Four participants were excluded due to SCR “lability” (non-responders). This refers to subjects who show spontaneous fluctuations of SCR in the absence of specific stimulation (Nonspecific Skin Conductance Response (NS-SCR) or slow SCR habituation (< 0.01 μS) (Raskin & Prokasy, 1973). According to Venables and Mitchell (Venables & Mitchell, 1996), approximately 25% of the normal population are SCR labile. The subjects’ lability was determined by visually tracking the real-time data during the experiment. Two further participants with more than 50% bad trials (movement artifacts and electrode artifacts) were excluded during the data pre-processing.

The final analysed data therefore are based on 25 retained participants (14 females; age mean=27.44 ± 9.76, age range 18 - 45).

#### Auditory stimuli

The auditory stimuli consisted of 15 s long pre-recorded single-sourced acoustic ‘scenes’ downloaded from ‘freesound.org’ and ‘sound-effects.bbcrewind.co.uk’ in two categories of ‘mechanical sounds’ and ‘nature sounds’. In each category, we had three distinctive sound scenarios: highway, jackhammer, and chainsaw for mechanical sounds and waterfall, birds chirping, and crickets for nature sounds. Of each scenario, four different exemplars were used. These stimuli were selected based on the International Affective Digitized Sounds database list – expanded version of the second edition (IADS - 2; (Lang & Bradley, 2007)). The IADS-E auditory stimuli (*N* = 935) are classified by their semantic categories, and the dataset includes the arousal and valence ratings collected from Japanese students (*N* = 207) (Yang et al., 2018). However, since the auditory stimuli in the IADS - E were short (1.5 s), we adopted the same stimuli (with the same label such as jackhammer in the mechanical sound category) with longer duration from other sources.

The objective of this experiment was to employ a diverse corpus of sounds that effectively encompassed each category of nature and mechanical sounds and were widely distributed across the valence and arousal spectra. For both natural and mechanical sound categories, the 0.25 (Q1), 0.5 (Q2), and 0.75 (Q3) quantiles were determined based on the arousal (nature ranging from 2.58 to 7.83; mechanical ranging from 2.18 to 8.58) and the valence scale (nature ranging from 1.82 to 8.25; mechanical ranging from 1.27 to 7.09). Subsequently, the sound exemplars whose mean arousal and valence values matched with Q1, Q2, and Q3 were methodically selected, including a waterfall (arousal 3.9, valence 6.6), highway (arousal 3.8, valence 5.6), crickets (arousal 5.2, valence 5), a chainsaw (arousal 5.4, valence 4.2), bird chirping (arousal 6.5, valence 3.7) and a jackhammer (arousal 7, valence 2.8), representing nature and mechanical sounds, respectively. This was done to guarantee that the chosen sounds would represent each category comprehensively, covering a broad spectrum of arousal and valence levels.

The sounds were normalized at three loudness levels: ∼ 10 sones (low), 20 sones (medium), and 30 sones (loud), making a total of 72 trials. The stimuli were normalized by ArtemiS SUITE HEAD acoustics software version 10.7 and measured in the laboratory by Head Acoustics SQobold with a BHS II headset. None of the auditory stimuli were above the hazardous threshold of 85 dB SPL (Costa et al., 2022).

Auditory stimuli were delivered to the participants through 12 coaxial loudspeakers (Genelec 8030A) placed to follow an imagined sphere (r=1.5 m) in three rows around a participant on the floor, 1.5 m height and 3.0 m height. The principle used to deliver 2-channel stereo recordings to the speaker array was based on routing the channel 1 to 6 speakers on the left side and the channel 2 to 6 speakers on the right side where the 4 middle speakers were assigned in a way to keep the balance of 2 speakers per channel on each height. Stimulus presentation was controlled with Psychtoolbox (Psychophysics Toolbox version 3; (Brainard, 1997) on MATLAB (version 2019a) (MATLAB, 2019)). The inter-stimulus interval varied between 30-60 seconds (randomly; in steps of 5 s).

### Loudness discrimination task

Prior to the main experiment, we conducted a short loudness discrimination task on participants to ensure the three loudness levels used were distinguishable. The participants were presented with one block which consisted of twelve pairs of stimuli selected from all three loudness levels. Of the twelve pairs, three pairs contained stimuli with the same loudness levels (low vs. low, medium vs. medium, and loud vs. loud), three pairs of low vs. medium, three pairs of low vs. loud, and three pairs of medium vs. loud. Participants were instructed to press a designated key on a provided keyboard to indicate the louder sound and to not react when sounds were presented at the same loudness level. The trials were randomized across participants **(Figure 1)**.

**Figure 1.**
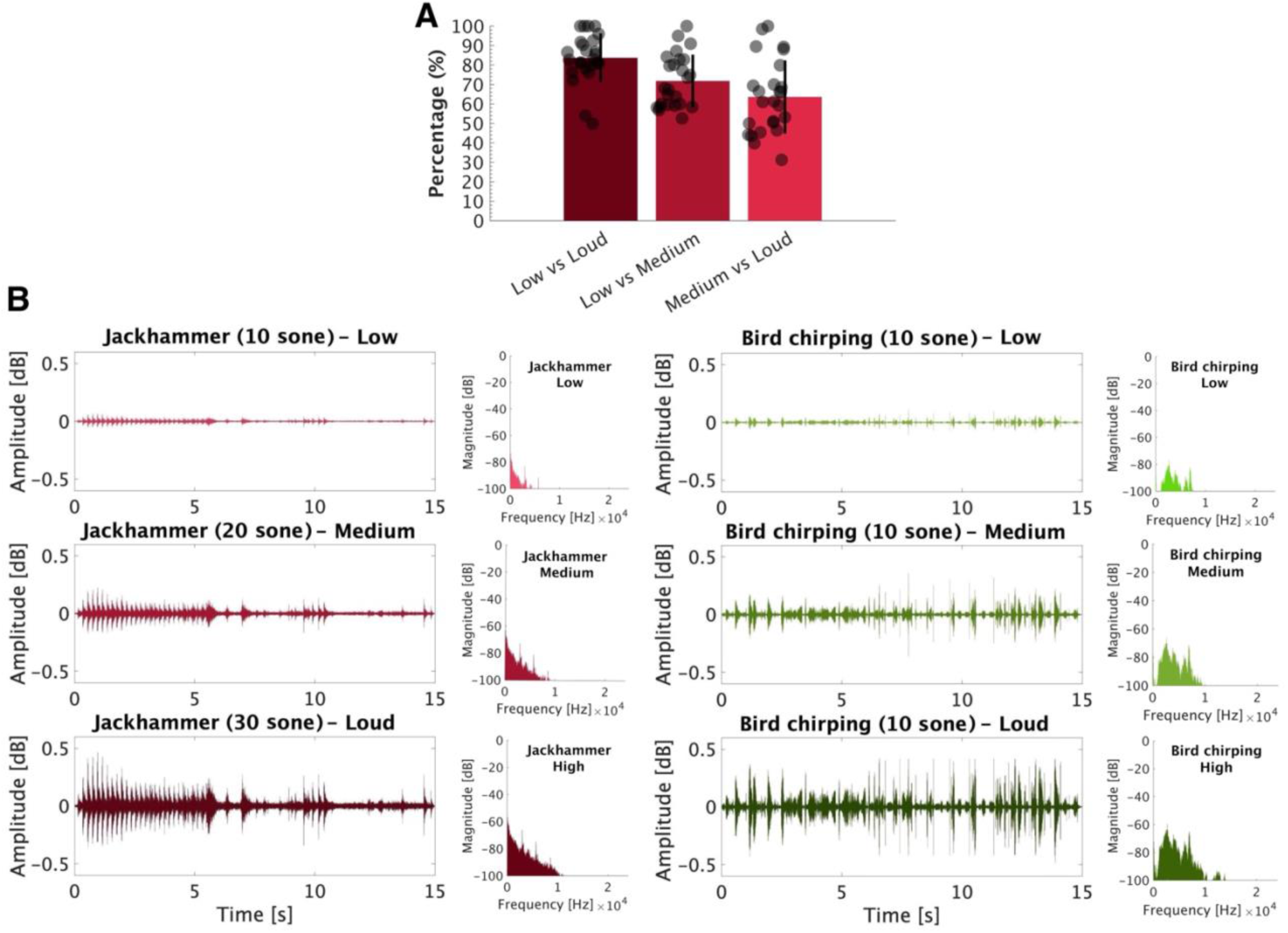
A) Loudness discrimination task. Error bars (SD) illustrate the variability in performance. B) Exemplars of mechanical (Jackhammer) and nature (Birds chirping) sounds at three loudness levels: low, medium, and loud in the time and frequency domains.

#### Procedure

The study consisted of two separate parts of ‘passive’ and ‘active’ listening tasks, followed by debriefing. To control for undesirable environmental factors such as noise interference, and temperature and to ensure that the SCR fluctuations were induced by the sound stimuli, we included 6 incidental trials (silence conditions (15 s)) in which no stimulus was presented to the participants. The silence conditions were spread randomly across blocks with one incidental trial in each block.

### Passive listening task

The participants sat in front of a monitor (23.8-in. BenQ 2480 Full HD with the WQHD resolution (2560 x 1440 pixels) in a dimly lit and acoustically shielded room. They were instructed to sit still and to continuously fixate on a changing colour fixation cross (∼2*2 cm) presented in the centre of the screen against a light grey background with a distance of ∼0.5 meters while being presented with the sounds. To assess the level of participants’ attention to the stimuli, they were directed to engage in a task where they were required to promptly press a space bar, located on a provided keyboard, as soon as the colour of the fixation cross changed (every ∼2-6 seconds). For an accurate recording of attention, a keypress was classified as a ‘hit’ only if it occurred within a time frame of less than 1.5 s following the change of the fixation cross colour. Auditory stimuli were presented randomly with interstimulus intervals of 30-60 s.

#### Skin Conductance Response (SCR)

The SCR was measured, using Empatica4 (CE Cert. No. 1876/MDD (93/ 42/EEC Directive, Medical Device class 2a - FCC CFR 47 Part 15b) which is an ambulatory device, normally wrapped around the wrist. Its reliability is comparable to clinical devices in appropriate circumstances (McCarthy et al., 2016). Since palmar and plantar areas showed to have stronger SCR and the wrist sweat glands may be primarily thermoregulatory in their functioning (Boucsein, 2012; Dawson et al., 2007), we attached the sensors of the device to the index and middle fingers of the participants’ non-dominant hands. The participants were instructed to minimize their movement during the experiment. SCR real-time data (via SCR sensor) and participants movement (via 3-axis accelerometer sensor) were continuously monitored and recorded by Empatica4-manager software (version 2.0.1.5023) at a sampling rate of 4 Hz. Each block started with 120 s SCR baseline measure to get a subtle signal and after each block, the data were automatically transferred to the Ematica4 secure cloud platform which was exported from the cloud for analysis.

To minimize fatigue, the participants were given breaks of 2 – 3 mins between each block and a long 5 – 10 mins break between the passive and the active listening tasks. At the end of the passive task, the Empatica4 was detached from the participants’ fingers. The SCR was measured only during the passive listening task.

### Active listening task

Subsequent to the recording of SCR during the passive listening task, an active listening task was conducted. The active listening task involved the random presentation of the same auditory sequence (15 s), with varying interstimulus intervals (30-60 s). Participants were instructed to rate each stimulus using 8 perceptual attributes, ranging from 0 (min) to 100 (max) during the given intervals by pressing the left and right navigation keys to move the cursor along a slide bar. All perceptual attributes were presented simultaneously on the same page, with their order counterbalanced across trials for each participant. For each trial, participants were allotted a maximum of 25 s (∼ 3 s for each attribute) to register their responses. If no response was recorded within this time frame, the trial was automatically advanced to the subsequent trial. All 8 perceptual attributes appeared after each sound stimulus for rating.

#### Perceptual attributes

The assessment of pleasantness and eventfulness was done by using an adapted version of ISO/TS 12913-3:2019, based on the Swedish Soundscape Quality Protocol (SSQP; 41) (ISO, 12913-3: 2019). It includes a question ‘To what extent do you agree/disagree that the present sound is …’. Using a continuous scale ranging from ‘0’ as the minimum to ‘100’ as the maximum, the participants evaluated the quality of the acoustic environment using eight adjectives: pleasant, chaotic, vibrant, uneventful, calm, annoying, eventful, and monotonous. Then, eight adjectives (PA) collapse into 2-dimentional coordinates plotted with continuous values between –100 to 100 for ‘pleasantness’ on the x-axis and ‘eventfulness’ on the y-axis. These dimensions were calculated as shown in formulas 1 and 2:

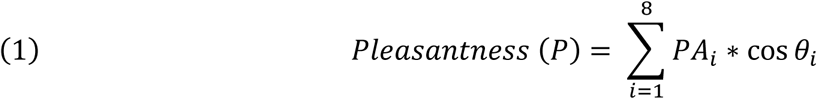

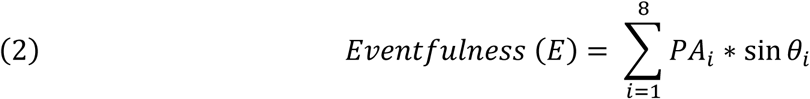

Where PA_1_ = pleasant, θ1 = 0°; PA_2_ = vibrant, θ2 = 45°; PA_3_ = eventful, θ3 = 90°; PA_4_ = chaotic, θ4 = 135°; PA_5_ = annoying, θ5 = 180°; PA_6_ = monotonous, θ6 = 225°; PA_7_ = uneventful, θ7 = 270°; PA_8_ = calm, θ8 = 315°.

The perceptual attributes as a measuring tool for the soundscape have been validated across several populations (Aletta et al., 2024).

#### SCR Pre-processing

The raw signal (recorded at 4Hz and 0.001-100 µS) in response to auditory stimuli was pre-processed and analysed by Continuous Decomposition Analysis (CDA), using the Ledalab toolbox (v. 3.2.2) MATLAB (version 2019a) (MATLAB, 2019)). Prior to the CDA, data were visually inspected (blinded to trial type), and movement artefacts were manually corrected by using spline interpolation. Then the CDA was carried out in four steps including deconvolution of SC data, estimation of tonic activity, estimation of phasic activity, and optimization.

These steps were initially performed for predefined parameters (τ_1_ = 1, τ_2_ = 3.75) to determine model fit. Subsequently, to enhance the goodness of the model, the parameters were optimized by re-applying these four steps.

Deconvolution of SC data: SC results from the convolution of sudomotor nerve bursts with the impulse response function (IRF), which describes the course of the impulse response over time. This process produces phasic and tonic components. We conducted deconvolution to reverse this process, allowing for the separation of phasic and tonic activity.
Estimation of tonic activity: Tonic electrodermal activity can occur without phasic activity (Boucsein, 2012), but the slow recovery of SCRs can obscure it. To estimate tonic activity, phasic response overlap was minimized by reducing its time constant, and then the time intervals between phasic impulses were used to estimate tonic activity. Deconvolution amplifies noise, so the tonic driver was smoothed using Gaussian convolution (σ = 0.2 s). Then peaks were detected by finding zero crossings in the first derivative, with significant peaks defined by a local maximum differing by δ ≥ 0.001 μS from adjacent minima. Non-overlapping sections estimated the tonic driver, which was interpolated using a cubic spline along a 10-s grid size. Finally, tonic SC activity was reconstructed by convolving the tonic driver with the IRF.
Estimation of phasic activity: The phasic driver was obtained by subtracting the tonic driver from the total driver signal, resulting in a signal with a near-zero baseline and positive deflections, capturing the time-constrained nature of phasic activity.
Optimization: The IRF can vary based on individual skin characteristics. Thus, our initial parameters (τ1 and τ2) were optimized based on criteria that evaluate the model. First, it was important for the phasic driver to display clear, short bursts of activity that quickly returned to zero between these bursts. To check how clear these bursts were (indistinctness), the number of consecutive bursts that were above a set level (5% of the maximum value of the phasic driver) were counted and then divided by the sampling rate (4Hz). This resulted in the duration of the bursts in seconds. Next, they were standardized by squaring, summing, and dividing by the total duration of the data, resulting in a measure in square seconds per second (s²/s). Higher values indicate longer bursts above the threshold. Second, the negative values in the phasic driver should be as low as possible. So, negativity was measured by calculating the RMS of the negative portions of the phasic driver. Finally, it created a criterion (c = indistinctiveness + negativity. a) that included both measures. The negativity measure was multiplied by a factor of α = 6 s²/ (s μS) to guarantee both measures contribute similarly to the criterion (Benedek & Kaernbach, 2010). The parameters were optimized by minimizing the criterion (c). A low (c) score indicates a good model with a stable baseline and clear bursts of activity. The optimization was achieved using a gradient descent method (Snyman & Wilke, 2005), which adjusted parameters to improve the criterion until no further significant improvements can be made (for more information see (Benedek & Kaernbach, 2010)).

After the CDA, the *CDA.SCR* from each trial was epoched from stimulus onset (*t_0_*) to 15 s post-onset (corresponding to the stimulus duration). CDA.SCR is the Ledalab metric that refers to the average phasic driver within the response window but does not rely on traditional SCR amplitude calculations [µS]. For each trial, baseline correction was applied by subtracting the mean SCR over the pre-onset interval (5 s pre-onset). Then data were averaged across all trials for each participant in each condition. The minimum threshold for SCR responses was set to 0.01µS.

For time series analysis, after the CDA, the *PhasicData* (phasic activity) from each trial were epoched from 5 s pre-stimulus to 25 s post-offset. For each trial, as in the previous instance, baseline correction was implemented by subtracting the mean SCR measured during the pre-onset interval. Then the data for each participant in each block were normalized. To do this, the mean and standard deviation (SD) across all baseline samples (5 s pre-onset interval) in each block were calculated and used to z-score normalize all data points (all epochs, all conditions) in the block. For each participant, SCR was time-domain averaged across all epochs of each condition to produce a single time series per condition.

#### Statistical analysis

Statistical analysis was conducted in MATLAB (version 2019a) (MATLAB, 2019)), and R statistical software (version 4.0.3). The *p-*value was a priori set to *p* < 0.05 for all analyses. If applicable, Greenhouse-Geisser correction for multiple comparisons was applied to all post hoc (Games-Howell) analyses conducted after ANOVA tests.

#### Time series statistical analysis

A non-parametric bootstrap-based analysis (Efron & Tibshirani, 1994) was used to evaluate time interval differences in SCR across conditions (‘silence’, ‘low‘,’ medium’, and ‘high’) and (‘nature’ and ‘mechanical’). For each participant, time series differences between conditions were computed and subjected to bootstrap resampling (1000 iterations with replacement). Statistical significance at each time point was determined by evaluating whether the proportion of bootstrap iterations exceeding (or falling below) zero surpassed the 95% confidence threshold (*p* < 0.05). Any significant differences observed during the pre-stimulus interval were attributed to noise. No difference was observed during the pre-stimulus interval.

## Results

### Incidental task performance is not modulated by loudness or sound category

For the incidental task (colour change detection), performance (d’) was computed using hit (HR) and false alarm rates [d’ = z (HR) – z (false alarms)]. A space bar press was considered a hit if it occurred within 1.5 s following the change in colour of the fixation cross. If HR or false alarms were at the ceiling, a standard correction was applied (Hautus, 1995). A parametric analysis (one-way ANOVA) with a Greenhouse-Geisser correction was performed to compare the conditions with factors of loudness level (10 sones ‘low’, 20 sones ‘mid’, and 30 sones ‘loud’). The performance measure revealed no difference between low (mean = 3.66, SEM = 0.04), mid (mean = 3.65, SEM = 0.04), and loud (mean = 3.63, SEM = 0.04) conditions F (2, 72) = 0.19, *p* = 0.83, indicating that performance on the incidental task was not affected by stimulus loudness.

Reaction times (RTs) were analysed from each HR with a two-way repeated-measures ANOVA (factors of loudness and category) with a Greenhouse-Geisser correction. The results revealed no significant simple main effect of loudness levels (F (1.83, 170.44) = 0.71, *p* = 0.48, η² = 0.008) or sound categories (F (1, 93) = 0.93, *p* = 0.34, η² = 0.01) on RTs and no (F (1.82, 169.63) = 0.01, *p* = 0.98, η² < 0.001). These results suggest that participants succeeded in maintaining their attention during the task regardless of loudness levels or sound categories.

### SCR is modulated by loudness but not sound category

The SCR was computed as the average conductance value over the 15-s-long stimulus presentation and evaluated with two-way repeated-measures ANOVA with a Greenhouse-Geisser correction, with factors of loudness level (‘silence’, ‘low’, ‘medium’, and ‘loud’) and sound category (‘nature, and ‘mechanical’). The results revealed a significant main effect of loudness, F (1.84,44.34) = 7.33, *p* = 0.002, η² = 0.23, indicating that the induced SCR varies across loudness levels. In contrast, there was no main effect of sound category, F (1,24) = 1.79, *p* = 0.19, η² = 0.07, confirming that SCR is strongly driven by the loudness levels rather than sound categories. Similarly, no interaction between loudness levels and sound categories was observed, F (1.69,40.78) = 0.84, *p* = 0.42, η² = 0.23 **(Figure 2)**.

**Figure 2.**
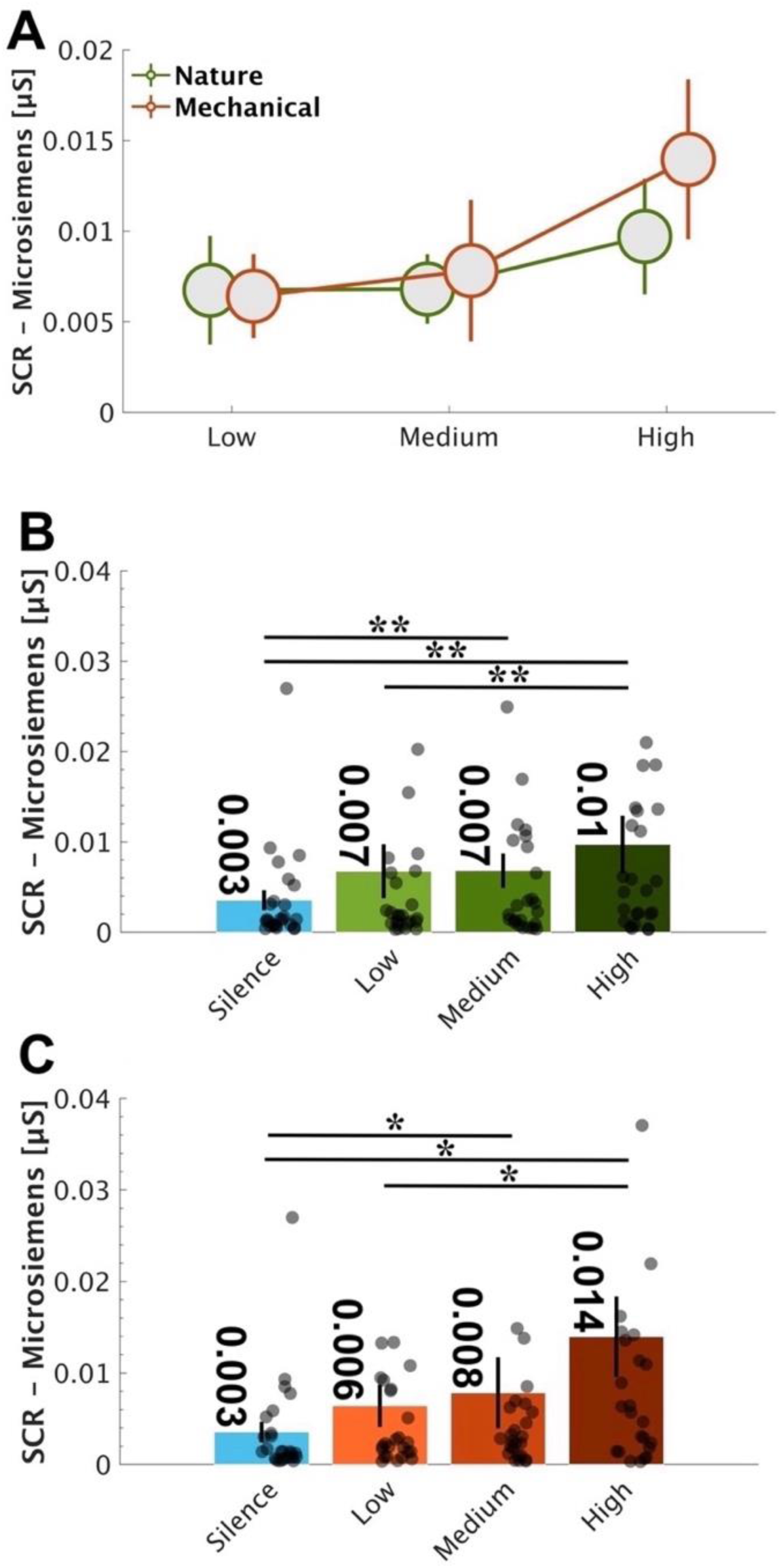
A) The SCR of all participants (*N* = 25). SCR was averaged across all trials and all participants. Error bars are ±1 SEM. B) The SCR of all participants in the nature and C) in the mechanical sound categories. Gray circles indicate individual data. Error bars are ±1 SEM. SCR measures were significantly modulated by increasing the loudness of nature and mechanical sounds (*** < 0.001, **< 0.01, *< 0.05).

**Figure 2B** plots the averaged SCR of all participants in the nature sound category and the mechanical sound **(Figure 2C)** category across loudness levels. A one-way ANOVA measure with a Greenhouse-Geisser correction was conducted with loudness level (‘silence’, ‘low’, ‘medium’, and ‘loud’) as factors across sound categories. To ensure that the elicited SCR was a result of stimulus presentation rather than NS-SCR, silence conditions were incorporated into the analysis. SCR measures yielded a main effect of loudness, χ2 = 4.92, *p* = 0.02, η² = 0.017 in nature sound condition. The post hoc test (Games-Howell test) demonstrated significant differences between loudness levels for silence (medium *p =* 0.005, loud *p =* 0.008), low (loud *p =* 0.003), medium (silence *p =* 0.005), and loud (silence *p =* 0.008, low *p =* 0.003). The differences between loudness levels were not significant including for silence (low *p =* 0.124), low (silence *p =* 0.124, medium *p =* 0.965), medium (low *p =* 0.965, loud *p =* 0.095), and loud (medium *p =* 0.095) (6 comparisons). Similarly, in the mechanical sound category, the SCR measures showed a main effect of loudness, χ2 = 4.27, *p* = 0.02, η² = 0.251. The post hoc test (Games-Howell test) demonstrated significant differences between some loudness levels for silence (medium *p =* 0.039, loud *p =* 0.01), low (loud *p =* 0.042), medium (silence *p =* 0.039), and loud (silence *p =* 0.01, low *p =* 0.042). On the other hand, some differences between loudness levels were not significant for silence (low *p =* 0.153), low (silence *p =* 0.153, medium *p =* 0.416), medium (low *p =* 0.416, loud *p =* 0.124), and loud (medium *p =* 0.124) (6 comparisons).

Overall, the pattern is consistent with a monotonic increase in SCR with loudness.

The SCR time domain data are presented in **Figure 3**. All conditions exhibit a prototypical pattern of an abrupt decrease followed by a sharp increase in SCR at ∼2 sec post onset, followed by a peak at about ∼3 sec post onset, with amplitude varying in proportion to loudness level, and then a slow decrease.

**Figure 3.**
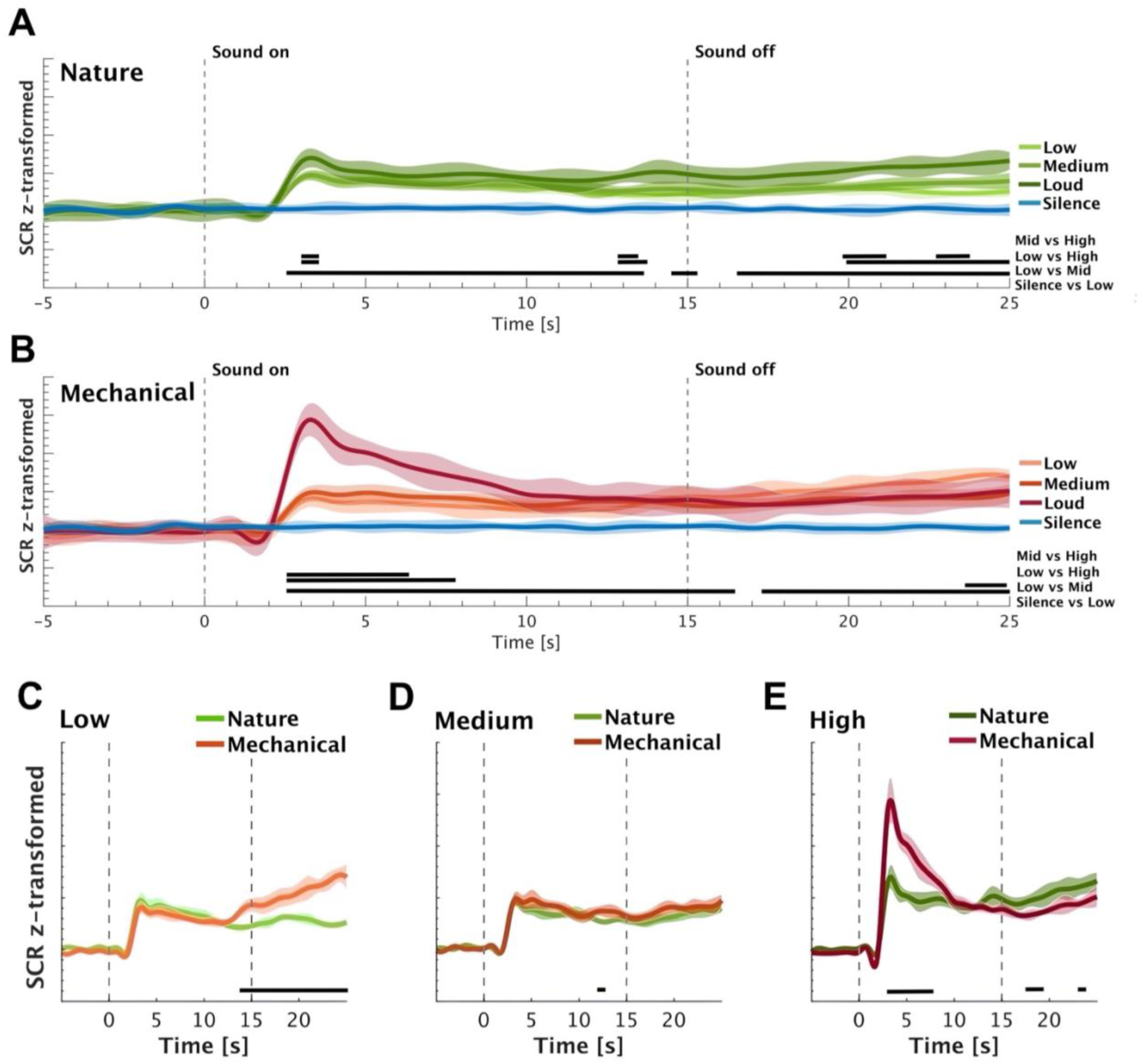
Comparison of the SCR across the conditions (‘silence’, ‘low’, medium’, and ‘loud’) in nature (A) and mechanical (B) sound categories. Panels C, D, and E illustrate the SCR in response to nature and mechanical sounds under low, medium, and high conditions, respectively. The shaded area shows ±1 SEM. The horizontal black lines represent the significant differences between conditions (*p <* 0.05).

To identify time intervals in which a given pair of conditions exhibited SCR differences, a non-parametric statistical analysis was used (bootstrap-based analysis) (demonstrated by the solid horizontal black lines in **Figure 3**). Especially following onset, a clear gradation by loudness was observed for both stimulus types. Loud mechanical sounds evoked the strongest response. We proceeded to compare nature and mechanical sounds separately for each loudness level. A difference was observed between loud nature and mechanical sounds, unfolding approximately 3 s after stimulus onset and persisting until ∼7 s post-stimulus presentation. This evidently suggests that loud mechanical sounds evoke stronger phasic activity relative to loud natural sounds, which may point to distinct characteristics of these sound categories. There was no significant difference between conditions at the ‘medium’ loudness level. However, at the ‘low’ loudness level, a notable divergence emerged late in the epoch, following sound offset: SCR to mechanical sounds exhibited an increase compared to those elicited by nature sounds. This pattern may suggest a delayed, slow-unfolding impact of mechanical sounds, even at low loudness levels. Given the incidental nature of this finding, however, we will not explore it further.

To gain a more comprehensive understanding of SCRs, we further analysed SCR velocity, which indicates the *speed* at which the SCR reaches its maximum following sound onset. The velocity of SCR was quantified as the peak derivative during the SCR rise-time for each trial per participant, capturing trial-specific dynamics. These trial-level velocities were then averaged within each participant to obtain a participant-specific mean velocity profile for each condition (e.g., low nature, high mechanical). Finally, these participant-level means were averaged across all participants within each condition to yield the group-level mean velocity, with SEM calculated to reflect inter-participant variability **(Figure 4).** For nature sounds **(Figure 4A)**, the velocity responses show a small, transient peak around 2 s post-stimulus onset, followed by a return to baseline. The magnitude of the response appears relatively consistent across conditions, with minimal difference in peak amplitude of loud nature sound. The data from mechanical sounds **(Figure 4B)** reveals a peak at approximately the same time point but with a greater difference between conditions. The SCR velocity in response to loud mechanical sounds elicits the largest peak, while the low and medium mechanical sounds show more attenuated responses. Direct comparisons within loudness conditions reveal, consistently with the previous analysis that high-loudness mechanical sounds are associated with higher SCR velocity than nature sounds

**Figure 4.**
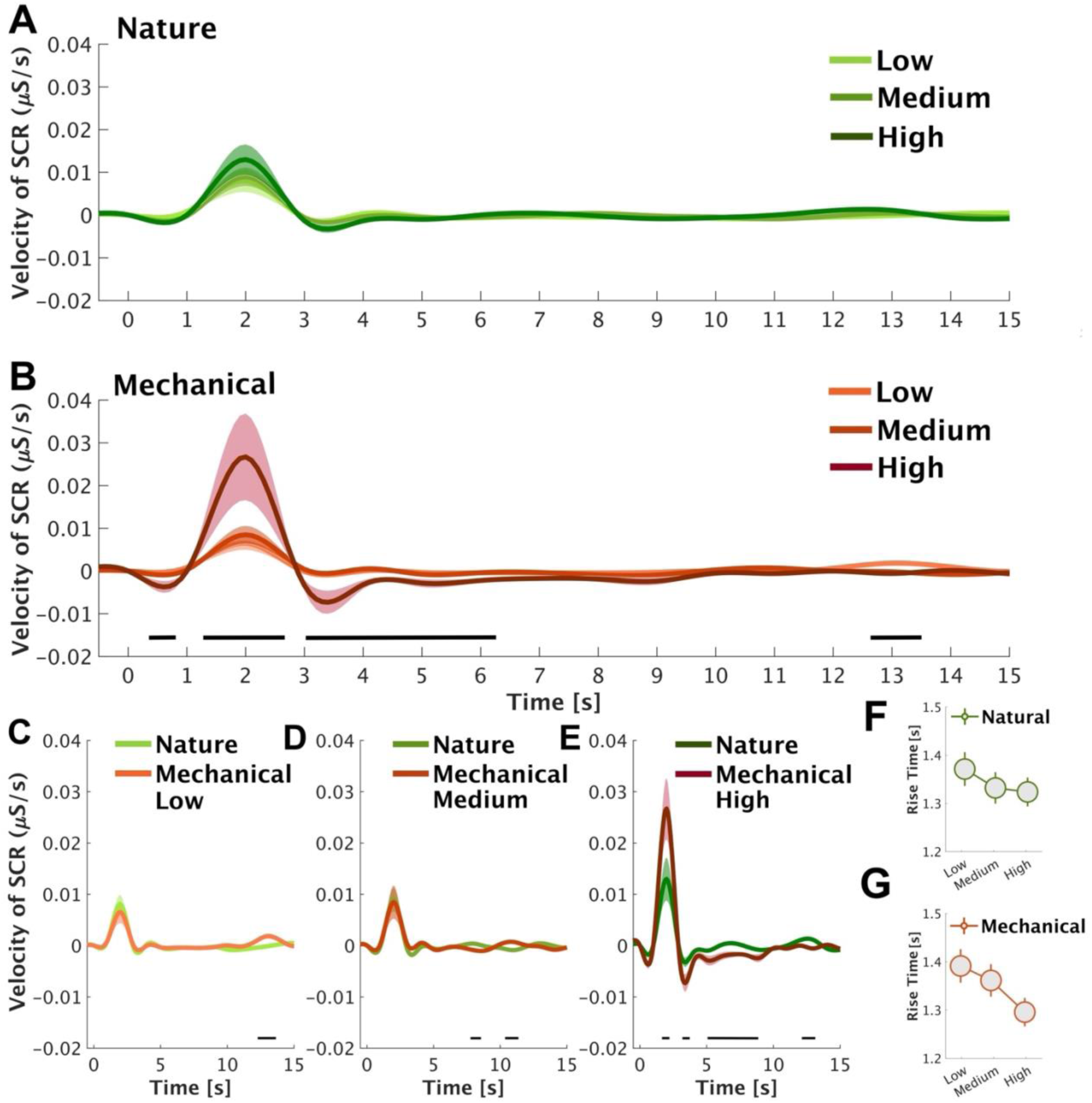
SCR velocity across loudness (‘silence’, ‘low’, medium’, and ‘loud’) (A & B) and sound category (‘nature’ and ‘mechanical’) (C, D, and E) conditions. The shaded area shows ±1 SEM. The black horizontal bars indicate time windows of significant differences. Panels F and G show SCR rise-time across loudness conditions.

In addition, we analysed the SCR rise-time. The SCR rise-time is identified as the point where the phasic component of the SC signal begins to increase following stimulus presentation, typically determined by a consistent upward deflection exceeding a predefined threshold (e.g., 0.01 µS). The peak is the maximum amplitude reached after onset, representing the highest conductance value before the response starts to decline. We tallied the SCR rise-time as the difference between the peak time and the SCR onset time, providing a measure of the *duration* at which the SCR reaches its maximum following initiation (Boucsein, 2012; Dawson et al., 2007; Venables & Christie, 1980). The SCR rise-time was computed for all conditions (‘low’, ‘medium’, and ‘loud’) across nature and mechanical sounds in all subjects. We ran two one-way ANOVAs which revealed no difference between loudness levels in nature F (2,72) = 0.5, *p* = 0.6, η² = 0.002, and mechanical F (2,72) = 1.55, *p* = 0.22, η² = 0.005 sound categories. No post hoc analysis was conducted **(Figure 4F and 4G)**.

### Pleasantness and eventfulness variance is driven by sound category

**Figure 5A** plots averaged soundscape pleasantness across all participants (*N* = 25). We evaluated soundscape pleasantness using repeated-measures analysis (two-way repeated ANOVA) with a Greenhouse-Geisser correction with factors of loudness level (‘low’, ‘medium’, and ‘loud’) and sound category (‘nature’ and ‘mechanical’). The analysis showed no main effect of loudness F (2,47.82) = 4.32, *p* = 0.19, η² = 0.15, indicating no difference in pleasantness between the loudness levels. The main effect of the sound category yielded a significant difference in soundscape pleasantness between nature and mechanical sounds, F (1,24) = 69.68, *p <* 0.001, η² = 0.74, with nature sounds judged as significantly more pleasant. No significant interaction was found between loudness and sound category (F (1.98,47.64) = 0.53, *p =* 0.59, η² = 0.02).

**Figure 5.**
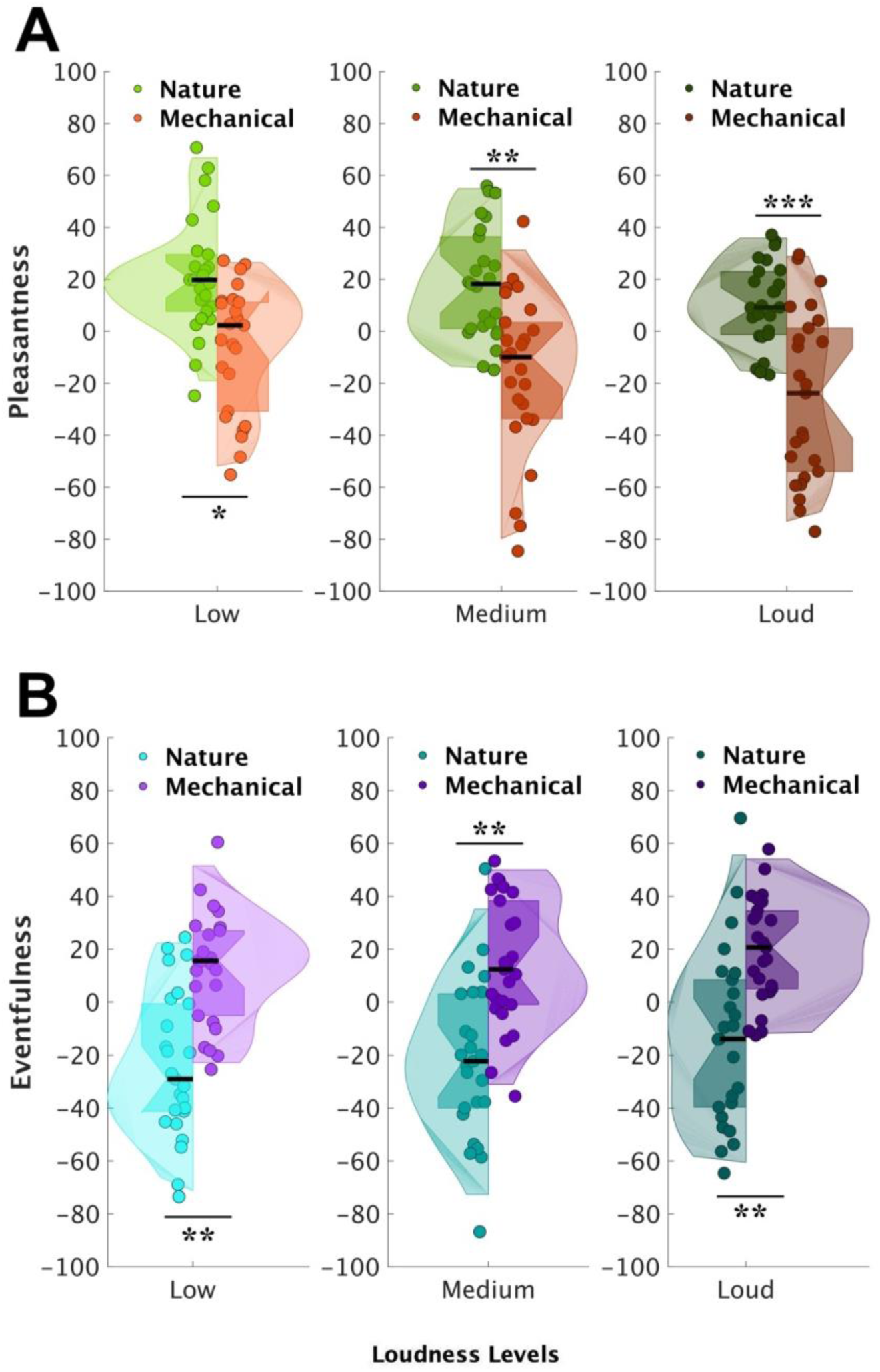
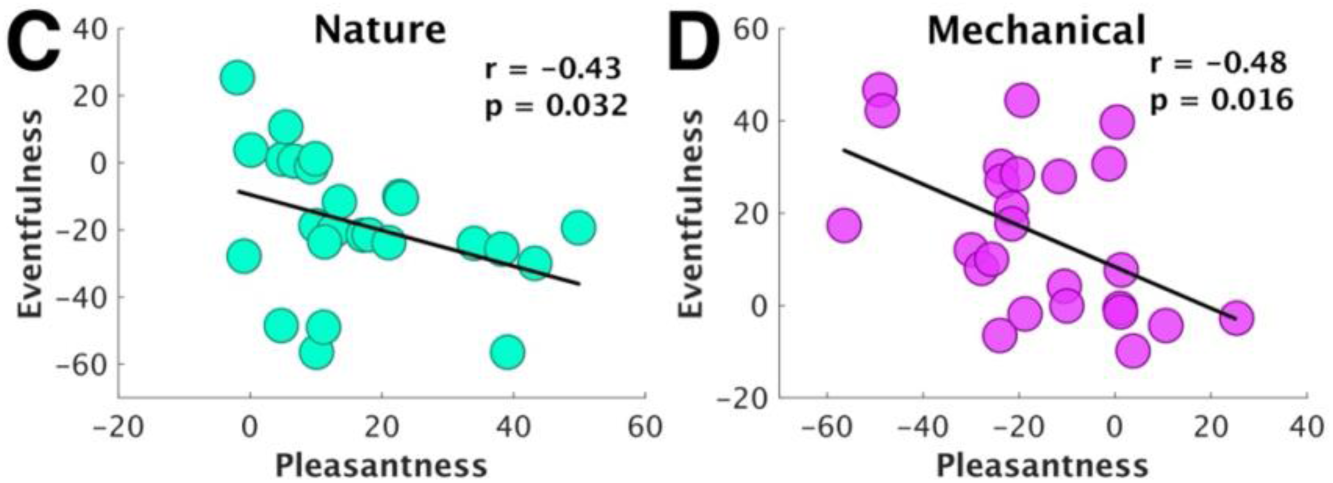
Soundscape pleasantness (A) and eventfulness (B) in response to nature and mechanical sounds across loudness levels (*N* = 25). The solid circles indicate the individual data. The black horizontal lines represent the mean. The upper end of the thick vertical rectangle shows mean + SD, and the lower end shows mean – SD. The bottom end of the thick rectangle illustrates the 1^st^ quartile, q (0.25), and the top end illustrates the 3^rd^ quartile, q (0.75). The bottom part of the kernel density shows the 1^st^ percentile, q (0.01) and the top part demonstrates the 99^th^ percentile, q (0.99). Panels C and D show correlations (Spearman) between soundscape pleasantness and eventfulness across nature and mechanical sound categories (*** < 0.001, **< 0.01, *< 0.05).

**Figure 5B** shows soundscape eventfulness across all participants (*N* = 25), evaluated with the same measure as soundscape pleasantness (two-way repeated-measures ANOVA with a Greenhouse-Geisser correction). Like the soundscape pleasantness, there was no main effect of loudness F (1.95,47.02) = 1.76, *p* = 0.18, η² = 0.07, whereas the main effect of the sound category was significant, revealing a difference in soundscape eventfulness between nature and mechanical sounds, with the latter being judged as significantly more “eventful” F (1,24) = 46.22 *p<* 0.001, η² = 0.66. No interaction was observed between loudness and sound category F (1.98,47.51) = 0.09, *p =* 0.92, η² = 0.004.

As has been reported previously (Erfanian et al., 2021; Mitchell et al., 2021), we observed a moderate negative correlation between pleasantness and eventfulness for both nature and mechanical sounds **(Figure 5C and 5D)**.

### Pleasantness and eventfulness do not correlate with SCR amplitude

Spearman correlation was employed to investigate potential links between soundscape pleasantness and eventfulness and the SCR amplitude, separately for nature and mechanical sound categories (**Figure 6A, B, C, D)**. The analysis revealed no significant correlations between soundscape pleasantness or eventfulness and the SCR amplitude across the two sound categories.

**Figure 6.**
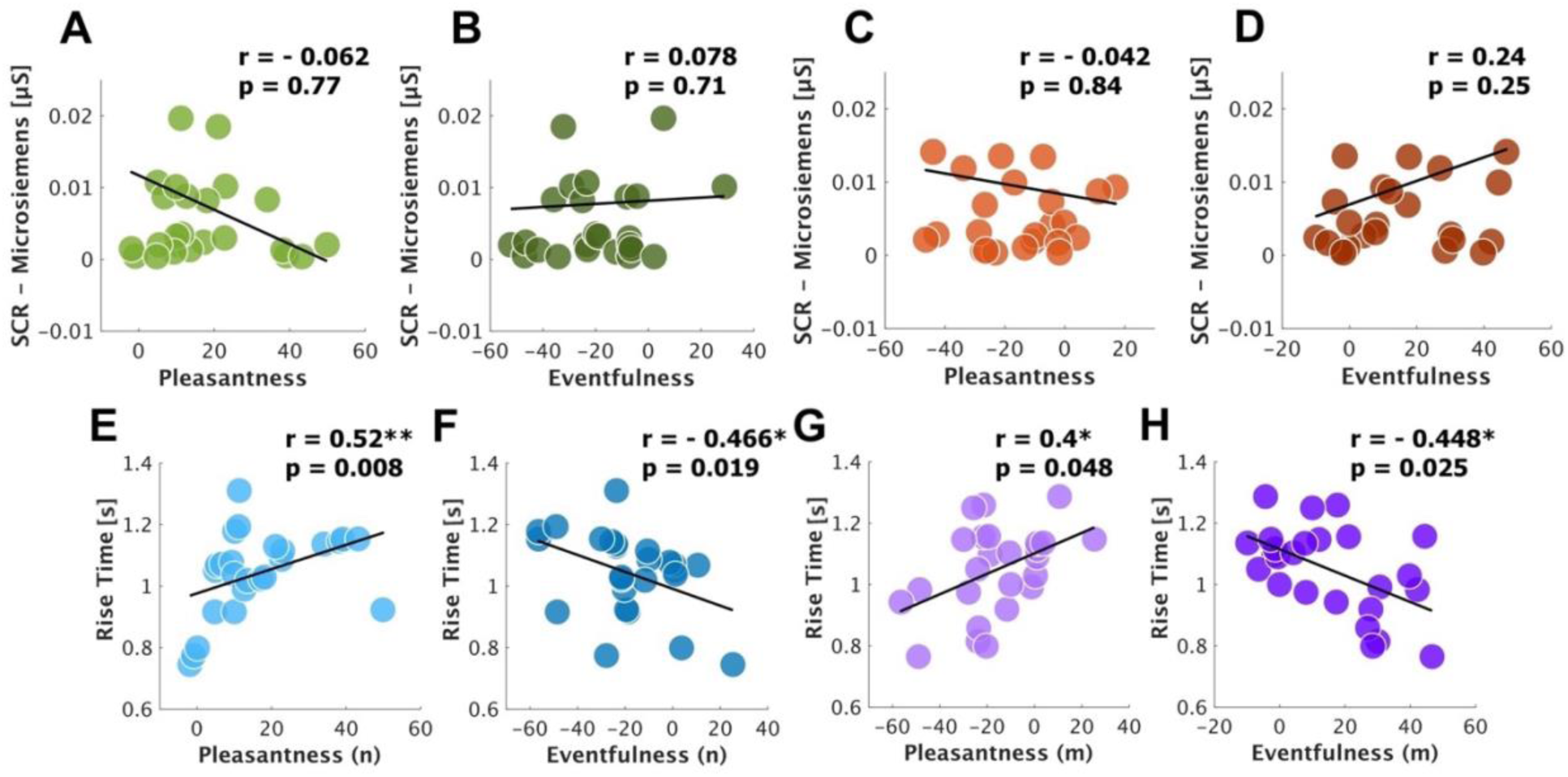
Correlation (Spearman) between soundscape pleasantness and eventfulness and the SCR amplitude and the SCR rise-time in seconds across nature (green and blue) and mechanical (orange and purple) sound categories (*N =* 25). Each circle represents individual data, and the diagonal line is the line of best fit, expressing the (degree of) relationships between the factors (*** < 0.001, **< 0.01, *< 0.05).

### Pleasantness and eventfulness correlate with SCR rise-time

To investigate whether there were associations between the SCR rise-time and the soundscape pleasantness and eventfulness, we correlated (Spearman) these factors across nature and mechanical sounds (**Figure 6**). Significant positive correlations between the SCR rise-time and the soundscape pleasantness were observed in nature (r= 0.52, *p* = 0.008) (**Figure 6E**) and mechanical sounds (r= 0.4, *p* = 0.048) **(Figure 6G)**, suggesting that the sounds with higher pleasantness prompted slower SCR rise-time across both sound categories. Inversely, we found significant negative moderate correlations between the SCR rise-time and the soundscape eventfulness in nature (r = -0.466, *p* = 0.019) **(Figure 6F)** and mechanical sounds (r= -0.448, *p =* 0.025) **(Figure 6H)** which evidenced that sounds with higher levels of eventfulness tend to elicit faster SCR rise-time in nature and mechanical sounds. When controlling for eventfulness, partial correlation analyses revealed that the associations between SCR rise-time and pleasantness were no longer significant in both nature (r = -0.011, *p* = 0.961) and mechanical sounds (r = 0.110, *p* = 0.609). Similarly, the correlations between SCR rise-time and eventfulness were non-significant when controlling for pleasantness (nature: r = 0.050, *p* = 0.817; mechanical: r = 0.260, *p* = 0.221). This is likely due to the opposing effects of pleasantness and eventfulness.

## Discussion

This study explored how loudness influences variance in soundscape pleasantness, eventfulness, and their associated SCR. It also investigated the distinct effects of natural versus mechanical sound categories on these affective and physiological measures. Findings reveal that loudness significantly modulates SCR, with time-series analysis showing that SCR differentiates physiological arousal between loud natural and mechanical sounds. No significant correlation emerged between SCR and subjective ratings of pleasantness or eventfulness, however, SCR rise-time showed significant associations with both pleasantness and eventfulness across sound categories. These results suggest that physiological arousal in response to soundscape is primarily driven by acoustic intensity, whereas perceptual qualities are more strongly tied to the nature of the sound source.

### Skin conductance response is influenced by loudness

The results demonstrated that SCR increased significantly as the loudness levels increased for both sound categories. Evidence has indicated that the SCR is highly sensitive to sound intensity (Bach, 2014; Bach et al., 2009; Bach et al., 2008; Björk, 1986; Boucsein, 2012; Bradley & Lang, 2000a; Cacioppo et al., 2007; Dawson et al., 2007; Gatti et al., 2018; Grings & Schell, 1969). For instance, Czepiel et al. (Czepiel et al., 2021) found that louder musical tones elicited greater SCR, while Bari et al. (Bari et al., 2024) reported that brief noise bursts (5 s) at higher intensity levels significantly increased SCR. Ellermeier and colleagues further corroborated these findings by demonstrating a direct correlation between the sound pressure level of environmental noise (e.g., vehicles passing by) and SCR magnitude (Ellermeier et al., 2020). Additionally, studies on background noise on ventilation equipment noise presented at levels ranging from 35 to 75 dBA SPL in 2 min blocks have shown that increasing auditory intensity results in greater electrodermal activity (Alvar & Francis, 2024).

Empirical data, in this regard, shows that rising sound intensity can be potentially perceived as salient warning cues or even a looming threat (Bach et al., 2008; Burow et al., 2005; LeDoux, 2022), and thereby prompts the recruitment of attentional and physiological resources to elicit adaptive responses. In a more precise manner, sound intensity effect on neural activity in the amygdala has been noted (Bach et al., 2008). The amygdala, in turn, plays a regulatory role in SNS activity through its projection to the hypothalamus. Importantly, the innervation of sweat glands is predominantly cholinergic and sympathetic in nature. This involves postganglionic fibres that originate from the sympathetic chain, as detailed by Shields et al. (Shields et al., 1987). This implies a compelling causal relationship between the intensity of sound and the episodic bursts of sympathetic nerve activity that eventually result in the manifestation of the SCR.

### Skin conductance is modulated by sound category only when it reaches high loudness levels

Our results indicate that SCR is predominantly modulated by stimulus loudness/intensity rather than the categorical nature of the auditory input. Studies using controlled auditory paradigms have demonstrated that sounds from distinct categories such as nature (e.g., rain, wind), human (e.g., vocalizations), and mechanical (e.g., alarms, machinery) elicit comparable SCR magnitudes when matched for intensity (Bradley & Lang, 2000a; Hedblom et al., 2017; Kong & Han, 2024). This finding is consistent with the established role of the SNS, mediated through its connection with the salience network (Seeley, 2019; Sturm et al., 2018; Xia et al., 2017), in responding primarily to the physical salience of sensory input as opposed to categorical or affective interpretation (Seeley, 2019). Moreover, neurophysiological models suggest that the amygdala (Cheng et al., 2007; Morris et al., 2001) and brainstem structures (Cacioppo et al., 2007; Dawson et al., 2007), via two relatively independent pathways leading to SCR generation (Boucsein, 2012; Edelberg, 1972), play a crucial role in mediating autonomic responses to auditory stimuli based on their acoustic properties rather than their semantic content (Dawson et al., 2007). The relative invariance of SCR to sound category highlights its role as a non-specific index of physiological arousal, mainly governed by low-level acoustic features rather than higher-order perceptual processing.

This observation, however, did not persist when contrasting the time-series data of loud nature sounds relative to mechanical sounds, as time-series analysis is sensitive to transient and rapidly fluctuating responses over time. This issue was discussed by (Bach & Friston, 2013), shedding light on the limitations of traditional operational approaches in SCR analysis which may overlook the temporal structure of physiological responses. Instead, they advocate for model-based methods that more accurately capture the underlying generative processes driving SCR dynamics. The differences in SCR between these conditions could be due to the dynamic interaction between loudness and frequency in real-world auditory perception, wherein changes in one parameter can affect the salience of the other (Neuhoff et al., 1999). Increasing loudness can enhance the prominence of specific frequency components and may even induce *pitch shifts* (Neuhoff et al., 2002). At higher loudness levels, acoustic properties inherent to mechanical sounds, such as their higher spectral content (Yang & Kang, 2013)) may become more pronounced. These psychoacoustic features are particularly salient and attention-grabbing, demanding greater cognitive resources and prioritizing threat detection. Consequently, mechanical sounds may elicit heightened autonomic arousal at higher levels, leading to stronger physiological responses (Storbeck & Clore, 2008).

### Soundscape pleasantness and eventfulness are mediated by sound category irrespective of loudness

The extant literature validates mechanical sounds are typically regarded as unpleasant, whereas nature sounds tend to elicit pleasantness. Each sound category possesses distinct acoustic characteristics (e.g., decibel level/intensity in mechanical sounds); (Nooralahiyan et al., 1998; Quaranta & Dimino, 2007) such that one, more than one or the interaction of these inherent acoustic characteristics modulate the pleasantness and eventfulness of the soundscape. Considering that loudness is an acoustic property strongly tied to unpleasantness (Mitchell et al., 2021), we controlled for loudness to examine the extent to which loudness, as an inherent acoustic feature, contributes to the soundscape pleasantness and eventfulness of nature versus mechanical sounds. The findings confirmed the previous work, evidencing that soundscape pleasantness evoked by nature sounds was significantly higher relative to mechanical sounds, even when matched for loudness level (e.g., 20 sones). In contrast, mechanical sounds were perceived as more eventful than nature sounds, even at equal loudness levels.

The present investigation is consistent with past research positing that nature-sound scenarios are generally rated as more pleasant and less eventful compared to their mechanical counterparts. This observation may be attributed to inherent acoustic features that are typically present in nature sounds including energy at low frequencies than at high (Voss & Clarke, 1978), and slow temporal modulations (Attias & Schreiner, 1997; Singh & Theunissen, 2003). Sounds containing elevated levels of energy at higher frequencies elicit aversive responses in listeners, while those with lower frequency content are more pleasant (Kumar et al., 2008; Patchett, 1979). Additionally, temporal modulation of sounds within the roughness range (30-150 Hz) has been shown to induce unpleasantness, aversion, and defence reactions (Arnal et al., 2019; Taffou et al., 2021). In this regard, mechanical sounds may predominantly inhered acoustic properties (e.g., roughness, sharpness, and loudness) that are tied to disgust, aversion, excitability, unpleasantness, and perceptual arousal. These findings imply that the perceptual attributes of the soundscape related to nature and mechanical sounds are unlikely to be accounted for by mere decibel level/intensity. Future research is warranted to determine and tease apart the degree of contribution of other psychoacoustic features, such as spectral content, within each sound category.

### Perceptual attributes demonstrate concordance with SCR rise-time but not is sustained phase

We observed no association between mean SCR (across the sound presentation epoch) and soundscape pleasantness and eventfulness, where eventfulness refers to the perceptual intensity and temporal dynamism of the auditory experience. We further investigated the SCR velocity and rise-time, which bears valuable information concerning the level of arousal (Dawson et al., 2007; Jindrová et al., 2020), thereby allowing for better discrimination of SCR to stimuli with varied degrees of arousal. The SCR rise-time is widely acknowledged to be inversely proportional to the magnitude of the gate current, (most SCR require a gate current of 0.1 - 50 mA (milliamp) to fire), and its build-up rate (Dawson et al., 2007). The results revealed that the sound category has a significant impact on SCR velocity and rise-time, exhibiting a negative and positive correlation with soundscape pleasantness and eventfulness, respectively. Expectedly, sounds with a positive valence, indicative of pleasantness, resulted in slower velocity and longer SCR rise-time, while those perceived as more eventful elicited faster velocity and shorter SCR rise-time, characterized by a steeper slope. These results are consistent with similar experiments in other modalities (Jindrová et al., 2020), which argued that stimuli characterized by high arousal and unpleasantness (negative valence and selected from the IAPS by Bradley & Lang (Bradley & Lang, 2000a)) elicited a faster/shorter SCR rise-time.

The velocity, a less commonly used SCR metric, and rise-time of SCR represent the speed and duration required for the skin electrical conductance to elevate from baseline to its peak level in response to a stimulus. These parameters are indices of the speed at which the SNS reacts. An SCR with a faster velocity and shorter rise-time signifies more rapid SNS activation, while a slower velocity and longer rise-time indicate slower activation. Nonetheless, factors such as the intensity of the stimulus and individual differences in SC levels may also impact these indices. The SNS is responsible for the ‘fight or flight’ response, an automatic reaction triggered by a perceived threat or danger. In such circumstances, the SNS must respond promptly to prepare the organism for a defensive reaction. Consequently, stimuli that possess greater perilous implications for the organism stimulate a more pronounced ‘fight or flight’ response, as demonstrated by a faster/shorter SCR velocity and rise-time, which enables the organism to effectively prepare for an evasive response to ward off the perceived danger (Storbeck & Clore, 2008).

## Conclusion

This study presents empirical evidence for the impact of loudness and sound category on soundscape perceptual and physiological attributes. Key findings indicate that while loudness levels modulate SCR, with SCR increasing as loudness rises, pleasantness and eventfulness remain unaffected. Conversely, the sound category (nature and mechanical) influences the pleasantness and eventfulness of the soundscape. The change in SCR does not correspond to the variance in pleasantness and eventfulness; however, SCR rise-time, which is inversely proportional to SCR amplitude, is associated with pleasantness and eventfulness. Collectively, this study provides validated insights into the acoustic properties that impact the affective dimensions of sound perception and the associated physiological substrates. These findings are advantageous for soundscape researchers, auditory neuroscientists, audiologists, and sound designers, who can use this knowledge to create healthier and more optimal acoustic environments by carefully considering relevant acoustic properties.

### CRediT Statement

ME conceptualized the study and developed the methodology with input from MC and JK. ME conducted data collection, performed formal analysis, and curated the data under the supervision of MC. TO assisted with laboratory preparation. Software development was handled by ME and TO. ME drafted the original manuscript, with editing conducted collaboratively and technical feedback by MC. MC and JK supervised the project. Additionally, MC and JK oversaw project administration and supported funding acquisition efforts.

## Acknowledgements

This project has received funding from the European Research Council (ERC) under the European Union Horizon 2020 research and innovation programme (grant agreement No. 740696).

## Data Availability Statement

The data reported in this manuscript alongside related information will be available on OSF upon publication.

## References

Aletta, F., Mitchell, A., Oberman, T., Kang, J., Khelil, S., Bouzir, T. A. K., Berkouk, D., Xie, H., Zhang, Y., & Zhang, R. (2024). Soundscape descriptors in eighteen languages: Translation and validation through listening experiments. Applied Acoustics, 224, 110109. 10.1016/j.apacoust.2024.110109

Aletta, F., & Torresin, S. (2023). Adoption of ISO/TS 12913-2: 2018 protocols for data collection from individuals in soundscape studies: An overview of the literature. Current Pollution Reports, 9(4), 710–723. 10.1007/s40726-023-00283-6

Alvar, A., & Francis, A. L. (2024). Effects of background noise on autonomic arousal (skin conductance level). JASA Express Lett, 4(1). 10.1121/10.0024272

Alvarsson, J. J., Wiens, S., & Nilsson, M. E. (2010). Stress recovery during exposure to nature sound and environmental noise. Int J Environ Res Public Health, 7(3), 1036–1046. 10.3390/ijerph7031036

Arnal, L. H., Kleinschmidt, A., Spinelli, L., Giraud, A.-L., & Mégevand, P. (2019). The rough sound of salience enhances aversion through neural synchronisation. Nature Communications, 10(1), 3671. 10.1038/s41467-019-11626-7

Attias, H., & Schreiner, C. (1997). Coding of naturalistic stimuli by auditory midbrain neurons. Advances in neural information processing systems, 10.

Axelsson, Nilsson, M. E., & Berglund, B. (2010). A principal components model of soundscape perception. J Acoust Soc Am, 128(5), 2836–2846. 10.1121/1.3493436

Bach, D. R. (2014). A head-to-head comparison of SCRalyze and Ledalab, two model-based methods for skin conductance analysis. Biological psychology, 103, 63–68. 10.1016/j.biopsycho.2014.08.006

Bach, D. R., Flandin, G., Friston, K. J., & Dolan, R. J. (2009). Time-series analysis for rapid event-related skin conductance responses. J Neurosci Methods, 184(2), 224–234. 10.1016/j.jneumeth.2009.08.005

Bach, D. R., & Friston, K. J. (2013). Model-based analysis of skin conductance responses: Towards causal models in psychophysiology. Psychophysiology, 50(1), 15–22. 10.1111/j.1469-8986.2012.01483.x

Bach, D. R., Schächinger, H., Neuhoff, J. G., Esposito, F., Salle, F. D., Lehmann, C., Herdener, M., Scheffler, K., & Seifritz, E. (2008). Rising sound intensity: an intrinsic warning cue activating the amygdala. Cerebral Cortex, 18(1), 145–150. 10.1093/cercor/bhm040

Bari, D. S., Aldosky, H. Y., Tronstad, C., & Martinsen, Ø. G. (2024). Disturbances in Electrodermal Activity Recordings Due to Different Noises in the Environment. Sensors, 24(16), 5434. 10.3390/s24165434

Benedek, M., & Kaernbach, C. (2010). A continuous measure of phasic electrodermal activity. Journal of neuroscience methods, 190(1), 80–91. 10.1016/j.jneumeth.2010.04.028

Björk, E. (1986). Laboratory annoyance and skin conductance responses to some natural sounds. Journal of sound and vibration, 109(2), 339–345.

Bones, O., Cox, T. J., & Davies, W. J. (2018). Sound categories: Category formation and evidence-based taxonomies. Frontiers in Psychology, 9, 331591. 10.3389/fpsyg.2018.01277

Boucsein, W. (2012). Electrodermal activity. Springer Science & Business Media.

Bradley, M. M., & Lang, P. J. (2000a). Affective reactions to acoustic stimuli. Psychophysiology, 37(2), 204–215. https://www.ncbi.nlm.nih.gov/pubmed/10731770

Bradley, M. M., & Lang, P. J. (2000b). Measuring emotion: Behavior, feeling, and physiology. Brainard, D. H. (1997). The Psychophysics Toolbox. Spat Vis, 10(4), 433-436. https://www.ncbi.nlm.nih.gov/pubmed/9176952

Burow, A., Day, H. E., & Campeau, S. (2005). A detailed characterization of loud noise stress: Intensity analysis of hypothalamo–pituitary–adrenocortical axis and brain activation. Brain research, 1062(1-2), 63–73. 10.1016/j.brainres.2005.09.031

Buxton, R. T., Pearson, A. L., Allou, C., Fristrup, K., & Wittemyer, G. (2021). A synthesis of health benefits of natural sounds and their distribution in national parks. Proceedings of the National Academy of Sciences, 118(14), e2013097118. 10.1073/pnas.2013097118

Cacioppo, J. T., Tassinary, L. G., & Berntson, G. (2007). Handbook of psychophysiology. Cambridge university press.

Carraturo, G., Kliuchko, M., & Brattico, E. (2024). Loud and unwanted: Individual differences in the tolerance for exposure to music. The Journal of the Acoustical Society of America, 155(5), 3274–3282. 10.1121/10.0025924

Cheng, D. T., Richards, J., & Helmstetter, F. J. (2007). Activity in the human amygdala corresponds to early, rather than late period autonomic responses to a signal for shock. Learn Mem, 14(7), 485–490. 10.1101/lm.632007

Costa, J. J., Al Kadir, S. T., Dhrubo, S. R., Wahid, M. F., Ratan, Z. A., & Roy, D. (2022). A Wearable Hearing Protection Device for Vehicle Drivers to Mitigate the Impact of Sound Pollution for Noisy Places in Bangladesh. 2022 International Conference on Innovations in Science, Engineering and Technology (ICISET),

Czepiel, A., Fink, L. K., Fink, L. T., Wald-Fuhrmann, M., Tröndle, M., & Merrill, J. (2021). Synchrony in the periphery: inter-subject correlation of physiological responses during live music concerts. Scientific reports, 11(1), 22457. 10.1038/s41598-021-00492-3

Dawson, M. E., Schell, A. M., & Filion, D. L. (2007). The electrodermal system. Handbook of psychophysiology, 2, 200–223.

Edelberg, R. (1972). Electrical activity of the skin: Its measurement and uses in psychophysiology. Handbook of psychophysiology, 367–418.

Efron, B., & Tibshirani, R. J. (1994). An introduction to the bootstrap. Chapman and Hall/CRC. 10.1201/9780429246593

Ellermeier, W., Kattner, F., Klippenstein, E., Kreis, M., & Marquis-Favre, C. (2020). Short-term noise annoyance and electrodermal response as a function of sound-pressure level, cognitive task load, and noise sensitivity. Noise Health, 22(105), 46–55. 10.4103/nah.NAH_47_19

Erfanian, M., Mitchell, A., Aletta, F., & Kang, J. (2021). Psychological well-being and demographic factors can mediate soundscape pleasantness and eventfulness: A large sample study. Journal of Environmental Psychology, 77, 101660. 10.1016/j.jenvp.2021.101660

Erfanian, M., Mitchell, A. J., Kang, J., & Aletta, F. (2019). The Psychophysiological Implications of Soundscape: A Systematic Review of Empirical Literature and a Research Agenda. Int J Environ Res Public Health, 16(19). 10.3390/ijerph16193533

Fruhholz, S., & Grandjean, D. (2013). Amygdala subregions differentially respond and rapidly adapt to threatening voices. Cortex, 49(5), 1394–1403. 10.1016/j.cortex.2012.08.003

Fruhholz, S., Trost, W., & Kotz, S. A. (2016). The sound of emotions-Towards a unifying neural network perspective of affective sound processing. Neurosci Biobehav Rev, 68, 96–110. 10.1016/j.neubiorev.2016.05.002

Gatti, E., Calzolari, E., Maggioni, E., & Obrist, M. (2018). Emotional ratings and skin conductance response to visual, auditory and haptic stimuli. Scientific data, 5(1), 1–12. 10.1038/sdata.2018.120

Gomez, P., & Danuser, B. (2004). Affective and physiological responses to environmental noises and music. International Journal of psychophysiology, 53(2), 91–103. 10.1016/j.ijpsycho.2004.02.002

Greco, A., Valenza, G., Citi, L., & Scilingo, E. P. (2016). Arousal and valence recognition of affective sounds based on electrodermal activity. IEEE Sensors Journal, 17(3), 716–725. 10.1109/JSEN.2016.262367

Grings, W. W., & Schell, A. M. (1969). Magnitude of electrodermal response to a standard stimulus as a function of intensity and proximity of a prior stimulus. J Comp Physiol Psychol, 67(1), 77–82. 10.1037/h0026651

Hautus, M. J. (1995). Corrections for extreme proportions and their biasing effects on estimated values of d′. Behavior research methods, instruments, & computers, 27, 46–51.

Hedblom, M., Gunnarsson, B., Iravani, B., Knez, I., Schaefer, M., Thorsson, P., & Lundstrom, J. N. (2019). Reduction of physiological stress by urban green space in a multisensory virtual experiment. Sci Rep, 9(1), 10113. 10.1038/s41598-019-46099-7

Hedblom, M., Knez, I., Ode Sang, A., & Gunnarsson, B. (2017). Evaluation of natural sounds in urban greenery: potential impact for urban nature preservation. R Soc Open Sci, 4(2), 170037. 10.1098/rsos.170037

Hume, K., & Ahtamad, M. (2013). Physiological responses to and subjective estimates of soundscape elements. Applied Acoustics, 74(2), 275–281. 10.1016/j.apacoust.2011.10.009

ISO12913-1. (2014). Acoustics—Soundscape—Part 1: Definition and Conceptual Framework. In: International Organization for Standardization Geneva.

ISO, T. (12913-3: 2019). Acoustics—Soundscape Part 3: Data Analysis. . ISO: Geneva, Switzerland.

Jindrová, M., Kocourek, M., & Telenský, P. (2020). Skin conductance rise time and amplitude discern between different degrees of emotional arousal induced by affective pictures presented on a computer screen. BioRxiv, 2020.2005. 2012.090829.

Jo, H. I., & Jeon, J. Y. (2020). Compatibility of data collection protocol in ISO 12913-2 for urban soundscape assessment. Forum Acusticum,

Kang, J., Aletta, F., Oberman, T., Erfanian, M., Kachlicka, M., Lionello, M., & Mitchell, A. (2019). Towards soundscape indices. Proceedings of the international congress on acoustics

Kong, P. R., & Han, K. T. (2024). Psychological and physiological effects of soundscapes: A systematic review of 25 experiments in the English and Chinese literature. Sci Total Environ, 929, 172197. 10.1016/j.scitotenv.2024.172197

Kumar, S., Forster, H. M., Bailey, P., & Griffiths, T. D. (2008). Mapping unpleasantness of sounds to their auditory representation. J Acoust Soc Am, 124(6), 3810–3817. 10.1121/1.3006380

Lang, P., & Bradley, M. M. (2007). The International Affective Picture System (IAPS) in the study of emotion and attention. Handbook of emotion elicitation and assessment, 29, 70–73.

Lang, P. J., & Bradley, M. M. (2010). Emotion and the motivational brain. Biological psychology, 84(3), 437–450. 10.1016/j.biopsycho.2009.10.007

LeDoux, J. E. (2022). As soon as there was life, there was danger: the deep history of survival behaviours and the shallower history of consciousness. Philosophical Transactions of the Royal Society B, 377(1844), 20210292. 10.1098/rstb.2021.0292

Li, Z., & Kang, J. (2019). Sensitivity analysis of changes in human physiological indicators observed in soundscapes. Landscape and Urban Planning, 190, 103593. 10.1016/j.landurbplan.2019.103593

MATLAB. (2019). Release, Statistics Toolbox, The MathWorks, Inc., Natick, Massachusetts, United States. In: ed.

McCarthy, C., Pradhan, N., Redpath, C., & Adler, A. (2016). Validation of the Empatica E4 wristband. 2016 IEEE EMBS international student conference (ISC),

McDermott, J. H. (2012). Auditory preferences and aesthetics: Music, voices, and everyday sounds. In Neuroscience of preference and choice (pp. 227–256). Elsevier. 10.1016/B978-0-12-381431-9.00020-6

Medvedev, O., Shepherd, D., & Hautus, M. J. (2015). The restorative potential of soundscapes: A physiological investigation. Applied Acoustics, 96, 20–26. 10.1016/j.apacoust.2015.03.004

Mitchell, A., Oberman, T., Aletta, F., Kachlicka, M., Lionello, M., Erfanian, M., & Kang, J. (2021). Investigating urban soundscapes of the COVID-19 lockdown: A predictive soundscape modeling approach. J Acoust Soc Am, 150(6), 4474. 10.1121/10.0008928

Morris, J. S., Buchel, C., & Dolan, R. J. (2001). Parallel neural responses in amygdala subregions and sensory cortex during implicit fear conditioning. Neuroimage, *13*(6 Pt 1), 1044-1052. 10.1006/nimg.2000.0721

Neuhoff, J. G., Kramer, G., & Wayand, J. (2002). Pitch and loudness interact in auditory displays: can the data get lost in the map? J Exp Psychol Appl, 8(1), 17–25. 10.1037//1076-898x.8.1.17

Neuhoff, J. G., McBeath, M. K., & Wanzie, W. C. (1999). Dynamic frequency change influences loudness perception: a central, analytic process. J Exp Psychol Hum Percept Perform, 25(4), 1050–1059. 10.1037//0096-1523.25.4.1050

Nooralahiyan, A., Kirby, H. R., & McKeown, D. (1998). Vehicle classification by acoustic signature. Mathematical and Computer Modelling, 27(9-11), 205–214.

Oszczapinska, U., Heller, L. M., Jang, S., & Nance, B. (2024). Ecological sound loudness in environmental sound representations. JASA Express Letters, 4(2). 10.1121/10.0024995

Patchett, R. F. (1979). Human sound frequency preferences. Perceptual and motor skills, 49(1), 324–326. 10.2466/pms.1979.49.1.324

Quaranta, V., & Dimino, I. (2007). Experimental training and validation of a system for aircraft acoustic signature identification. Journal of aircraft, 44(4), 1196–1204. 10.1121/10.0024995

Raskin, D. C., & Prokasy, W. F. (1973). Electrodermal Activity in Psychological Research. Academic Press, 417.

Salamon, J., Jacoby, C., & Bello, J. P. (2014). A dataset and taxonomy for urban sound research. Proceedings of the 22nd ACM international conference on Multimedia,

Schweiger, A., & Maltzman, I. (1985). Behavioural and electrodermal measures of lateralization for music perception in musicians and nonmusicians. Biological psychology, 20(2), 129–145. 10.1016/0301-0511(85)90021-3.

Seeley, W. W. (2019). The Salience Network: A Neural System for Perceiving and Responding to Homeostatic Demands. J Neurosci, 39(50), 9878–9882. 10.1523/JNEUROSCI.1138-17.2019

Shields, S. A., MacDowell, K. A., Fairchild, S. B., & Campbell, M. L. (1987). Is mediation of sweating cholinergic, adrenergic, or both? A comment on the literature. Psychophysiology, 24(3), 312–319. 10.1111/j.1469-8986.1987.tb00301.x

Singh, N. C., & Theunissen, F. E. (2003). Modulation spectra of natural sounds and ethological theories of auditory processing. The Journal of the Acoustical Society of America, 114(6), 3394–3411. 10.1121/1.1624067

Skagerstrand, Å., Köbler, S., & Stenfelt, S. (2017). Loudness and annoyance of disturbing sounds–perception by normal hearing subjects. International Journal of Audiology, 56(10), 775–783. 10.1080/14992027.2017.1321790

Snyman, J. A., & Wilke, D. N. (2005). Practical mathematical optimization (Vol. 97). Springer.

Storbeck, J., & Clore, G. L. (2008). Affective arousal as information: How affective arousal influences judgments, learning, and memory. Social and personality psychology compass, 2(5), 1824–1843. 10.1111/j.1751-9004.2008.00138.x

Sturm, V. E., Brown, J. A., Hua, A. Y., Lwi, S. J., Zhou, J., Kurth, F., Eickhoff, S. B., Rosen, H. J., Kramer, J. H., Miller, B. L., Levenson, R. W., & Seeley, W. W. (2018). Network Architecture Underlying Basal Autonomic Outflow: Evidence from Frontotemporal Dementia. J Neurosci, 38(42), 8943–8955. 10.1523/JNEUROSCI.0347-18.2018

Taffou, M., Suied, C., & Viaud-Delmon, I. (2021). Auditory roughness elicits defense reactions. Sci Rep, 11(1), 956. 10.1038/s41598-020-79767-0

Tarlao, C., Steele, D., & Guastavino, C. (2022). Assessing the ecological validity of soundscape reproduction in different laboratory settings. PLoS One, 17(6), e0270401. 10.1371/journal.pone.0270401

Tavano, A., & Poeppel, D. (2019). A division of labor between power and phase coherence in encoding attention to stimulus streams. Neuroimage, 193, 146–156. 10.1016/j.neuroimage.2019.03.018

Venables, P. H., & Christie, M. J. (1980). Electrodermal activity. Techniques in psychophysiology, 54(3).

Venables, P. H., & Mitchell, D. A. (1996). The effects of age, sex and time of testing on skin conductance activity. Biol Psychol, 43(2), 87–101. 10.1016/0301-0511(96)05183-6

Voss, R. F., & Clarke, J. (1978). ’’1/f noise’’in music: Music from 1/f noise. The Journal of the Acoustical Society of America, 63(1), 258–263. 10.1121/1.381721

Wallin, B. G. (1981). Sympathetic nerve activity underlying electrodermal and cardiovascular reactions in man. Psychophysiology, 18(4), 470–476. 10.1111/j.1469-8986.1981.tb02483.x

Xia, C., Touroutoglou, A., Quigley, K. S., Feldman Barrett, L., & Dickerson, B. C. (2017). Salience Network Connectivity Modulates Skin Conductance Responses in Predicting Arousal Experience. J Cogn Neurosci, 29(5), 827–836. 10.1162/jocn_a_01087

Yang, M., & Kang, J. (2013). Psychoacoustical evaluation of natural and urban sounds in soundscapes. J Acoust Soc Am, 134(1), 840–851. 10.1121/1.4807800

Yang, W., Makita, K., Nakao, T., Kanayama, N., Machizawa, M. G., Sasaoka, T., Sugata, A., Kobayashi, R., Hiramoto, R., & Yamawaki, S. (2018). Affective auditory stimulus database: An expanded version of the International Affective Digitized Sounds (IADS-E). Behavior Research Methods, 50, 1415–1429. 10.3758/s13428-018-1027-6

